## Supplemental Figures and Tables for "How inhibitory and excitatory inputs gate output of the inferior olive"

### Supplemental tables

| Source of the synaptic response | Type of stimulation | Latency of synaptic response (ms) (mean ± SD) | Half width V dep. (mean ± SD) | Half width V hyp. (mean ± SD) | Half width V reb. (mean ± SD) |
|---|---|---|---|---|---|
| CN | ChrimsonR | 14.35 ± 6.74 | -------- | 73.59 ± 14.97 | 79.74 ± 33.09 |
|  | GAD2Cre/H134R | 29.33 ± 10.08 | -------- | 77.89 ± 7.49 | 82.98 ± 17.79 |
| MDJ | Electrical | 0.57 ± 0.23 | 12.06 ± 3.06 | 78.93 ± 11.09 | 79.16 ± 14.98 |
| | Chronos | 2.26 ± 0.42 | 11.86 ± 5.03 | 80.90 ± 19.39 | 86.13 ± 12.96 |

**Supplemental Table 1. Kinetic properties of CN and MDJ synaptic responses using different stimulation approaches.** Latency and half width of hyperpolarization ($V_{hyp.}$) and rebound ($V_{reb.}$) synaptic components of CN synaptic responses evoked in mice transduced with ChrimsonR-tdTomato in the CN and GAD2Cre/Chr2-H134R-EYFP transgenic mice line, and latency and half width of depolarization ($V_{dep.}$), hyperpolarization ($V_{hyp.}$), and rebound ($V_{reb.}$) synaptic components of MDJ synaptic response evoked in mice transduced with Chronos in the MDJ and electrical stimulation.

| Stimulation | Slope ± SD | Intercept ± SD | $r^2$ | Number of cells |
|---|---|---|---|---|
| -100 ms | -0.49 ± 0.03 | -0.16 ± 0.02 | 0.69 | 6 |
| -50 ms | -0.41 ± 0.05 | -0.11 ± 0.02 | 0.44 | 6 |
| 0 ms | -0.56 ± 0.02 | 0.08 ± 0.01 | 0.88 | 5 |
| +30 ms | -0.53 ± 0.03 | 0.02 ± 0.01 | 0.78 | 4 |
| +50 ms | -0.51 ± 0.02 | 0.02 ± 0.01 | 0.74 | 6 |
| +70 ms | -0.45 ± 0.05 | 0.05 ± 0.02 | 0.5 | 5 |
| +100 ms | -0.48 ± 0.03 | 0.09 ± 0.01 | 0.71 | 5 |
| +150 ms | -0.69 ± 0.05 | 0.35 ± 0.03 | 0.61 | 7 |
| +200 ms | -0.51 ± 0.03 | 0.08 ± 0.01 | 0.74 | 4 |
| IPSP | -0.52 ± 0.01 | 0.07 ± 0.01 | 0.86 | 5 |
| MDJ Spike | -0.50 ± 0.02 | 0.09 ± 0.01 | 0.82 | 5 |
| CN spike | -0.54 ± 0.04 | 0.15 ± 0.02 | 0.78 | 8 |
| +150 ms Spike | -0.66 ± 0.04 | 0.33 ± 0.02 | 0.72 | 7 |

**Supplemental Table 2**. **PRC parameters of different stimulation paradigms.** Slope, Y-intercept, and $R^2$ of PRCs generated by dual (CN and MDJ afferents stimulation) stimulation at different time intervals (all of them evoking subthreshold synaptic responses except for '+150 ms spike' which evokes suprathreshold synaptic responses), CN afferent stimulation evoking subthreshold and suprathreshold synaptic responses ('IPSP' and 'CN spike', respectively), and MDJ afferent stimulation evoking suprathreshold synaptic responses ('MDJ spike').

| Uncorrected Fisher's LSD | Mean Diff. | 95.00% CI of diff. | P- value |
|---|---|---|---|
| + 150 ms vs. - 100 ms | -0.20 | -0.31 to -0.10 | 0.0001 |
| + 150 ms vs. - 50 ms | -0.28 | -0.39 to -0.18 | <0.0001 |
| + 150 ms vs. 0 ms | -0.13 | -0.24 to -0.022 | 0.018 |
| + 150 ms vs. +30 ms | -0.17 | -0.27 to -0.061 | 0.002 |
| + 150 ms vs. +50 ms | -0.18 | -0.28 to -0.089 | 0.0001 |
| + 150 ms vs. +70 ms | -0.24 | -0.34 to -0.14 | <0.0001 |
| + 150 ms vs. +100 ms | -0.22 | -0.32 to -0.11 | <0.0001 |
| + 150 ms vs. +200 ms | -0.19 | -0.29 to -0.082 | 0.0005 |
| + 150 ms vs. IPSP | -0.17 | -0.26 to -0.078 | 0.0003 |
| + 150 ms vs. MDJ spike | -0.19 | -0.29 to -0.096 | <0.0001 |
| + 150 ms vs. CN spike | -0.15 | -0.28 to -0.021 | 0.023 |
| + 150 ms vs. +150 ms spike | -0.029 | -0.14 to 0.077 | 0.59 |

**Supplemental Table 3. Statistics of PRC slopes of different stimulation paradigms.** Comparison of PRC slopes between +150 ms and the other time intervals (all of them evoking subthreshold synaptic responses except for '+150 ms spike' which evokes suprathreshold synaptic responses), CN afferent stimulation evoking subthreshold and suprathreshold synaptic responses ('IPSP' and 'CN spike', respectively) and MDJ afferent stimulation evoking suprathreshold synaptic responses ('MDJ spike'), using one-way ANOVA test followed by post-hoc uncorrected Fisher's LSD multiple comparison test.

| Uncorrected Fisher's LSD | Mean Diff. | 95.00% CI of diff. | P-value |
|---|---|---|---|
| + 150 ms vs. -100 ms | 0.52 | 0.46 to 0.58 | <0.0001 |
| + 150 ms vs. -50 ms | 0.46 | 0.41 to 0.52 | <0.0001 |
| + 150 ms vs. 0 ms | 0.27 | 0.21 to 0.33 | <0.0001 |
| + 150 ms vs. +30 ms | 0.33 | 0.27 to 0.39 | <0.0001 |
| + 150 ms vs. +50 ms | 0.33 | 0.28 to 0.38 | <0.0001 |
| + 150 ms vs. +70 ms | 0.30 | 0.24 to 0.36 | <0.0001 |
| + 150 ms vs. +100 ms | 0.26 | 0.20 to 0.32 | <0.0001 |
| + 150 ms vs. +200 ms | 0.26 | 0.21 to 0.32 | <0.0001 |
| + 150 ms vs. IPSP | 0.28 | 0.23 to 0.33 | <0.0001 |
| + 150 ms vs. MDJ spike | 0.26 | 0.20 to 0.31 | <0.0001 |
| + 150 ms vs. CN spike | 0.19 | 0.12 to 0.27 | <0.0001 |
| + 150 ms vs. +150 ms spike | 0.023 | -0.037 to 0.083 | 0.45 |

**Supplemental Table 4. Statistics of PRC Y-intercept.** Comparison of PRC Y-intercepts between +150 ms and the other time intervals (all of them evoking subthreshold synaptic responses except for '+150 ms spike' which evokes suprathreshold synaptic responses), CN afferent stimulation evoking subthreshold and suprathreshold synaptic responses ('IPSP' and 'CN spike', respectively) and MDJ afferent stimulation evoking suprathreshold synaptic responses ('MDJ spike') using one-way ANOVA test followed by post-hoc uncorrected Fisher's LSD multiple comparison test.

| **Stimulation** | **Intertrial phase jitter (Standard deviation of phase lags) pre-stimulus (ms)** | **Intertrial phase jitter (Standard deviation of phase lags) post-stimulus (ms)** | **Intertrial phase jitter change (Standard deviation of phase lags post stimulus – Standard deviation of phase lags pre-stimulus)** | **Number of cells** |
|---|---|---|---|---|
| **-100 ms** | **60.84** | **13.47** | **-47.37** | **10** |
| **-50 ms** | **58.39** | **19.86** | **-38.53** | **13** |
| **0 ms** | **56.87** | **14.84** | **-42.03** | **14** |
| **+30 ms** | **62.36** | **16.61** | **-45.74** | **17** |
| **+50 ms** | **64.84** | **36.48** | **-28.35** | **35** |
| **+70 ms** | **73.35** | **28** | **-45.34** | **17** |
| **+100 ms** | **57.51** | **18.13** | **-39.38** | **16** |
| **+150 ms** | **49.17** | **14.88** | **-34.98** | **10** |
| **+200 ms** | **52.61** | **10.62** | **-41.99** | **10** |
| **IPSP** | **59.51** | **20.35** | **-39.16** | **15** |
| **EPSP** | **56.47** | **44.97** | **-11.50** | **15** |
| **MDJ Spike** | **51.92** | **36.05** | **-9.29** | **8** |
| **+150 ms Spike** | **51.47** | **3.21** | **-48.26** | **7** |
| **CN spike** | **61.05** | **38.1** | **-22.95** | **7** |

**Supplemental Table 5. Intertrial phase jitter at different stimulation paradigms.** Intertrial phase jitter pre- and post-stimulus (Standard deviation of phase lags pre and post- stimulus, respectively) and intertrial phase jitter change (Standard deviation of phase lags post stimulus – Standard deviation of phase lags pre-stimulus) at different time intervals of stimulation (all of them evoking subthreshold synaptic responses except for '+150 ms spike' which evokes suprathreshold synaptic responses), CN afferent stimulation evoking subthreshold and suprathreshold synaptic responses ('IPSP' and 'CN spike', respectively) and MDJ afferent stimulation evoking subthreshold and suprathreshold synaptic responses ('EPSP' and 'MDJ spike', respectively).

| Time interval | Uncorrected Dunn's test | Rank sum diff. | P-value |
|---|---|---|---|
| **-100 ms** | **Δ Pre stim STO average vs. Post stim STO 1st peak** | **-276** | **<0.0001** |
| **-50 ms** | **ΔPre stim STO average vs. Post stim STO 1st peak** | **-346** | **<0.0001** |
| **0 ms** | **ΔPre stim STO average vs. Post stim STO 1st peak** | **-396** | **<0.0001** |
| **+30 ms** | **ΔPre stim STO average vs. Post stim STO 1st peak** | **-402** | **<0.0001** |
| **+ 50ms** | **ΔPre stim STO average vs. Post stim STO 1st peak** | **-452** | **<0.0001** |
| **+70 ms** | **ΔPre stim STO average vs. Post stim STO 1st peak** | **-53** | **0.07** |
| **+100 ms** | **ΔPre stim STO average vs. Post stim STO 1st peak** | **-440** | **<0.0001** |
| **+150 ms** | **ΔPre stim STO average vs. Post stim STO 1st peak** | **-307** | **<0.0001** |
| **+200 ms** | **ΔPre stim STO average vs. Post stim STO 1st peak** | **-367** | **<0.0001** |

**Supplemental Table 6. Statistics of amplitude of first cycle following stimulation at different time intervals.** Comparison between STO amplitude change of the first cycle (rebound component) following stimulation at different time intervals (all of them evoking subthreshold synaptic responses) and STO amplitude previous to stimulation using Friedman test followed by post-hoc uncorrected Dunn’s multiple comparison test.

| **Stimulation** | **STO amplitude change of 1st cycle post-stim ± SD (mV)** | **Number of cells** |
|---|---|---|
| **-100 ms** | **6.18 ± 4.98** | **10** |
| **-50 ms** | **5.53 ± 5.03** | **13** |
| **0 ms** | **4.80 ± 4.32** | **14** |
| **+30 ms** | **3.66 ± 2.98** | **16** |
| **+50 ms** | **1.83 ± 3.42** | **34** |
| **+70 ms** | **0.48 ± 2.59** | **17** |
| **+100 ms** | **6.48 ± 3.47** | **16** |
| **+150 ms** | **7.25 ± 3.45** | **10** |
| **+200 ms** | **4.08 ± 2.05** | **10** |
| **IPSP** | **4.82 ± 3.70** | **23** |
| **EPSP** | **1.52 ± 3.04** | **26** |
| **MDJ Spike** | **6.75 ± 4.26** | **10** |
| **CN spike** | **4.04 ± 3.19** | **6** |
| **+150 ms spike** | **8.67 ± 5.03** | **6** |

**Supplemental Table 7. STO amplitude change of first cycle following stimulation at different stimulation paradigms.** STO amplitude change of the first cycle (rebound component) following dual stimulation at different time intervals (all of them evoking subthreshold synaptic responses except for “+150 ms spike” which evokes suprathreshold synaptic responses),CN afferent stimulation evoking subthreshold and suprathreshold synaptic responses (‘IPSP’ and ‘CN spike’, respectively), and MDJ afferent stimulation evoking subthreshold and suprathreshold synaptic responses (‘EPSP’ and ‘MDJ spike’, respectively).

| Uncorrected Dunn's test | Mean rank diff. | P-value |
|---|---|---|
| **+150 ms vs. -100 ms** | **227.0** | **0.008** |
| **+150 ms vs. -50 ms** | **277.1** | **0.0005** |
| **+150 ms vs. 0 ms** | **359.1** | **<0.0001** |
| **+150 ms vs. +30 ms** | **522.2** | **<0.0001** |
| **+150 ms vs. +50 ms** | **787.1** | **<0.0001** |
| **+150 ms vs. +70 ms** | **1042** | **<0.0001** |
| **+150 ms vs. +100 ms** | **88.41** | **0.25** |
| **+150 ms vs. +200 ms** | **420.1** | **<0.0001** |
| **+150 ms vs. EPSP** | **867.1** | **<0.0001** |
| **+150 ms vs. IPSP** | **367.4** | **<0.0001** |
| **+150 ms vs. MDJ spike** | **110.1** | **0.19** |
| **+150 ms vs. CN spike** | **456.8** | **<0.0001** |
| **+150 ms vs. +150 ms spike** | **-37.74** | **0.68** |

**Supplemental Table 8. Statistics of the comparison between first cycle post-stimulus change evoked by +150 ms and other stimulation paradigms.** Comparison of first cycle post-stimulus change between the time interval of +150 ms and the other time intervals (all of them evoking subthreshold synaptic responses except for “+150 ms spike” which evokes suprathreshold synaptic responses), CN afferent stimulation evoking subthreshold and suprathreshold synaptic responses (‘IPSP’ and ‘CN spike’, respectively), and MDJ afferent stimulation evoking subthreshold and suprathreshold synaptic responses (‘EPSP’ and ‘MDJ spike’, respectively), using Kruskal-Wallis test followed by post-hoc uncorrected Dunn’s multiple comparison test.

| Stimulation | $P_{spike}$ ± SD | Number of cells |
|---|---|---|
| **-100 ms** | **0.16 ± 0.22** | **9** |
| **-50 ms** | **0.17 ± 0.26** | **9** |
| **0 ms** | **0.23 ± 0.39** | **7** |
| **+30 ms** | **0.05 ± 0.09** | **7** |
| **+50 ms** | **0.0 ± 0.0** | **9** |
| **+70 ms** | **0.0 ± 0.0** | **6** |
| **+100 ms** | **0.08 ± 0.21** | **8** |
| **+150 ms** | **0.57 ± 0.30** | **9** |
| **+200 ms** | **0.26 ± 0.22** | **7** |
| **IPSP** | **0.01 ± 0.03** | **7** |
| **EPSP** | **0.22 ± 0.28** | **7** |

**Supplemental Table 9**. **Spike probability at different stimulation paradigms.** Spike probability ($P_{spike}$) following dual stimulation at different time intervals and isolated CN and MDJ afferents stimulation ('IPSP' and 'MDJ', respectively).

| Uncorrected Fisher's LSD | Mean Diff. | 95.00% CI of diff. | P-value |
|---|---|---|---|
| +150 ms vs. -100 ms | 0.41 | 0.21 to 0.62 | 0.0002 |
| +150 ms vs. -50 ms | 0.4 | 0.19 to 0.61 | 0.0002 |
| +150 ms vs. 0ms | 0.34 | 0.13 to 0.54 | 0.0015 |
| +150 ms vs. +30 ms | 0.52 | 0.30 to 0.74 | <0.0001 |
| +150 ms vs. +50 ms | 0.58 | 0.37 to 0.78 | <0.0001 |
| +150 ms vs. +70 ms | 0.58 | 0.35 to 0.81 | <0.0001 |
| +150 ms vs. +100 ms | 0.49 | 0.28 to 0.70 | <0.0001 |
| +150 ms vs. +200 ms | 0.31 | 0.094 to 0.53 | 0.0057 |
| +150 ms vs. EPSPs only | 0.36 | 0.14 to 0.58 | 0.0018 |
| +150 ms vs. IPSP only | 0.56 | 0.34 to 0.78 | <0.0001 |

**Supplemental Table 10. Statistics of the comparison between spike probability evoked by +150 ms and other stimulation paradigms.** Comparison of spike probability ($P_{spike}$) of time interval of +150 ms and all the other time intervals and CN and MDJ isolated afferents stimulation ( 'IPSP' and 'EPSP', respectively) using one-way anova test followed by post-hoc uncorrected Fisher's LSD multiple comparison test.

#### Supplemental figure legends

**Supplemental Figure 1. MDJ evoked synaptic responses in the IO.** (**A**) Synaptic responses to increasing stimulus intensities using electrical stimulation. Arrows indicate how depolarizing ($V_{dep.}$), hyperpolarizing ($V_{hyp.}$) and rebound ($V_{reb.}$) synaptic components amplitudes were calculated. (**B**) $V_{dep.}$, $V_{hyp.}$ and $V_{reb.}$ amplitudes (mV) plotted as a function of stimulus intensity during electrical stimulation (mA), respectively. Open black circles represent individual trials. (**C**) Synaptic responses to increasing stimulus intensities using optical stimulation at a wavelength of 470nm in WT mice transduced with Chronos in the MDJ. (**D**)**,** $V_{dep.}$, $V_{hyp.}$ and $V_{reb.}$ amplitudes (mV) plotted as a function of stimulus intensity during optical stimulation at a wavelength of 470nm in WT mice transduced with Chronos in the MDJ, respectively. Open blue circles represent individual trials. (**E**) Example trace of synaptic response to MDJ inputs before and after the presence of picrotoxin (PTX, 100μM), CNQX (20 μM) and APV (50 μM). (**F**), $V_{dep.}$, $V_{hyp.}$ and $V_{reb.}$ normalized amplitude before and after the presence of CNQX, PTX and APV, respectively. Red lines indicate the mean and light gray lines indicate individual experiments.

**Supplemental Figure 2. PRC calculation.** (**A**) Period previous to stimulation ($T_0$) was calculated as the average of 4 periods preceding the stimuli. Period containing the synaptic responses caused by either single or dual stimuli ($T_1$) was calculated as the period from the peak cycle before stimulation to the peak cycle following the rebound peak caused by either single or dual stimulus. Time at which stimulation hit the STO ($T_s$) was calculated as the period from the peak cycle before stimulation to the onset of the first stimulus. (**B**) PRC curve in which changes in the period containing the synaptic responses with respect to the period previous to stimulation ($\Delta\Phi$) is plotted as function of the phase at which stimulus hit the STO ($\Phi$). Based in our convention shown in the formulas of panel A, positive $\Delta\Phi$ values indicate a phase advance (period containing the synaptic response is shorter than previous ones) whereas negative $\Delta\Phi$ values indicate a phase delay (period containing the synaptic response is longer than previous ones).

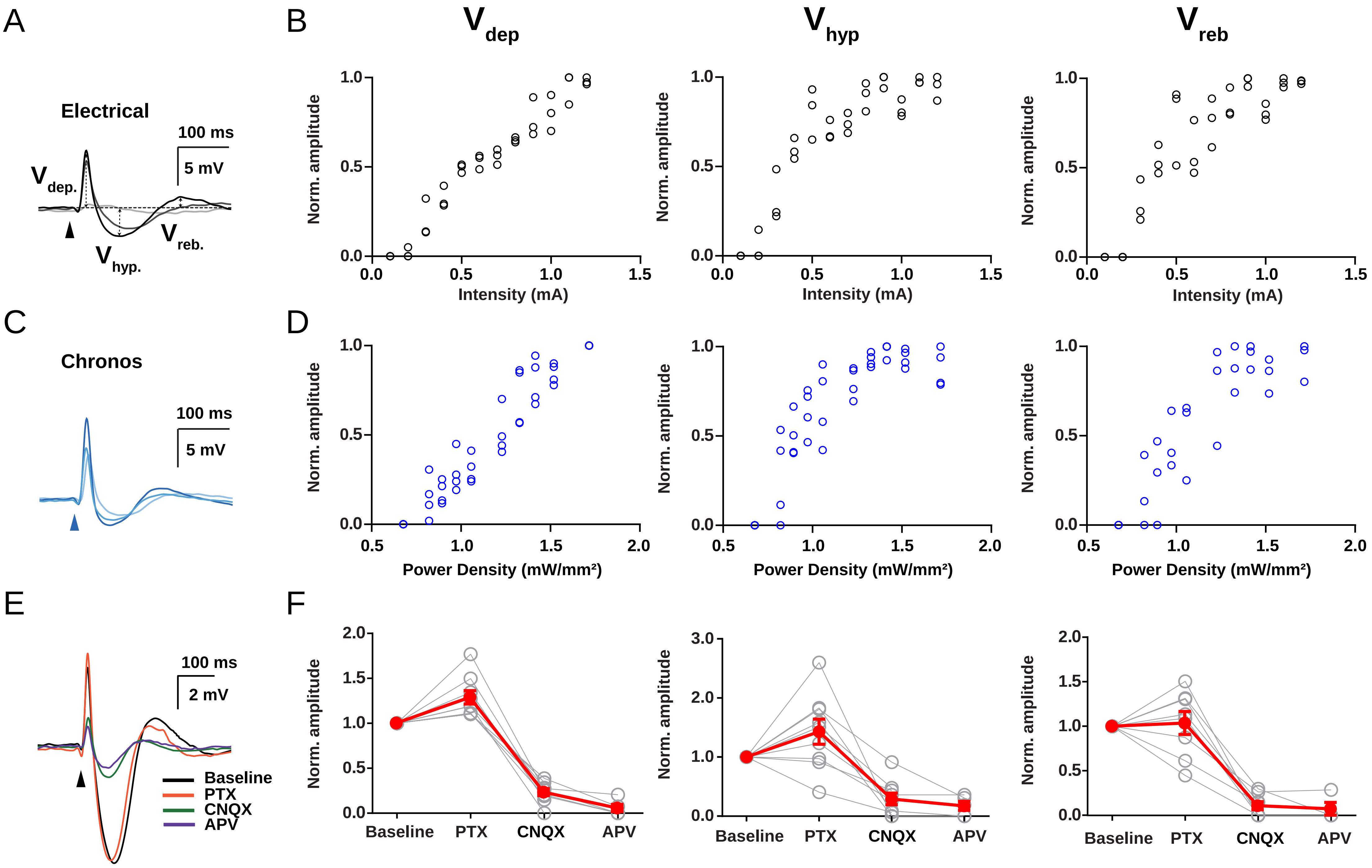


**Supplemental Figure 1**

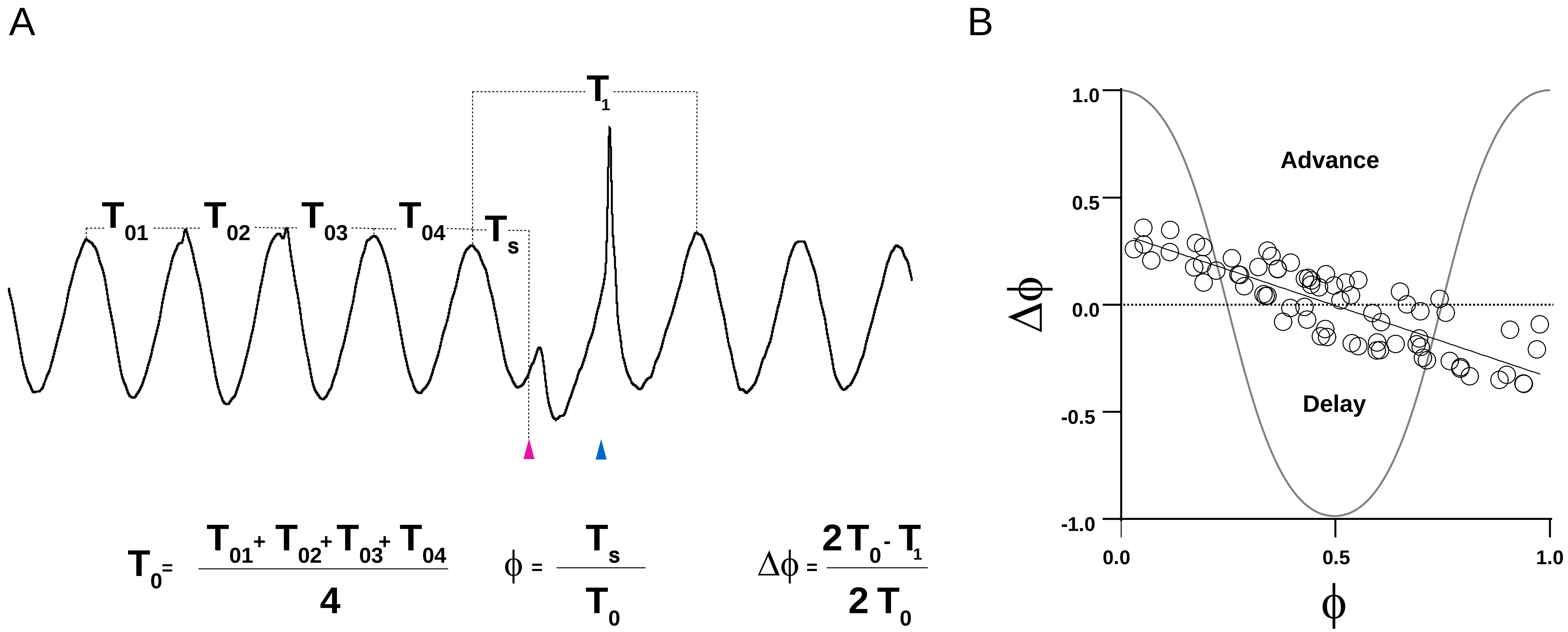


**Supplemental Figure 2**
